# *In vivo* de-amplification of a multi-resistance pseudo-compound transposon in Escherichia coli

**DOI:** 10.64898/2026.03.06.710075

**Authors:** Ralfh Pulmones, Sabrina J Moyo, Berhe Tesfay, Joshua Gidabayda, Museveni Justine, Iren Hoyland Lohr, Bjorn Blomberg, Simon Wagstaff, Nina Langeland, Adam P. Roberts

## Abstract

The rapid expansion of antimicrobial resistance (AMR) among Gram-negative pathogens presents a major clinical challenge, particularly in vulnerable populations such as infants. The dissemination of resistance is often mediated by mobile genetic elements (MGEs) that can mobilise antimicrobial resistance genes (ARGs) both within and between genomes. The insertion sequence (IS) IS*26*, is a MGE with the ability to replicate itself and associated ARGs, create translocatable units (TU), and produce tandem arrays of ARGs.

Here we present the 18-week *in vivo* evolution of a community-acquired multi-drug resistant (MDR) *Escherichia coli* colonising an infant gut, characterised by a de-amplification of an IS*26*-mediated tandem array of ARGs. The hybrid-assembled ancestral and descendant strain genomes show an evolutionary rate of 10.22 SNPs per genome per year. Independent analysis of the hybrid genome assembly, and of Oxford Nanopore Technologies (ONT) and Illumina read-mapping support the existence of at least five copies of a TU (Tn*3*-like(*tnpA*)-*tetR-tetA-yedA-*ΔTn*1721*(*tnpA*) *-*IS*26*-*aac(6’)-Ib-cr-bla_OXA-1_-*Δ*catB3*-IS*26*) in the ancestor and only one in the descendant. Despite de-amplification, no change in fitness (p = 0.275) and piperacillin-tazobactam susceptibility (TZP) was observed. In contrast, gentamicin susceptibility increased, in the absence of known associated mutations.

This study provides insight into IS*26*-mobility dynamics *in vivo* and their implications for AMR, within the rapidly changing environment of the neonatal gut.

## Introduction

Antimicrobial resistance (AMR) is a present and growing threat to global health. Among the most threatening antimicrobial resistant bacteria are extended-spectrum beta lactamase (ESBL)-carrying Gram-negative bacteria, acknowledged by the World Health Organisation as an urgent threat and a top priority for drug development and research (1). Mediating the dissemination of AMR are Mobile Genetic Elements (MGEs) which can mobilise both themselves and associated DNA, often including antimicrobial resistance genes (ARGs).

MGEs are genetic elements capable of excising and inserting within a genome and, in the case of mobilizable and conjugative MGEs, transferring between cells. These include transposons, insertion sequences (ISs) and plasmids. Transposons can typically transpose by one of two mechanisms: replicative and non-replicative transposition, although some can utilise both (2). Non-replicative transposition involves a transposase gene (*tnpA*) producing an enzyme which recognises its own flanking terminal inverted repeats and initiating excision (3). Most known transposons are non-replicative, but some, such as the Tn*3*-family of transposons, are capable of replicative transposition. In Tn*3* and some ISs, replicative transposition involves *tnpA*-mediated co-integration between the transposon and target DNA molecule (e.g., a chromosome or plasmid), which is dis-entangled by a site-specific resolvase (*tnpR*). This leaves a copy of the IS at the original site and the new target, and short target site duplications (TSDs) (4).

IS*26* canonically cannot transpose alone and requires a nearby indirectly oriented IS*26* to mobilise, as a composite transposon. These composite transposons typically use a replicative “copy-in” mechanism in which a co-integrate is formed between the composite transposon and the target molecule. IS*26* composite transposons must have indirectly-oriented IS*26* because the co-integrates they form must be resolved by *recA*-mediated homologous recombination, because IS*26* lacks a *tnpR* (5). However, a variant of replicative transposition, targeted conservative co-integration (TCC), was described for IS*26* by Harmer and Hall, which requires directly-oriented IS*26* to form a pseudo-compound transposon (PCT) (6). Through TCC, a translocatable unit (TU) comprising of IS*26* and cargo DNA can transpose adjacent to another directly oriented IS*26*. In turn, the TU can be copied and the process repeated, creating a tandem array of repeated TUs with no TSDs (7). This can increase the copy number of multiple ARGs simultaneously which can increase resistance phenotypes, or expand to newer antibiotic generations and combination therapies (8,9). Amplification of ARGs, and the spread of MDR, is especially concerning for vulnerable populations like infants.

In Tanzania, mortality from bloodstream infections (BSIs) of primarily ESBL-*Enterobacteriaceae* (ESBL-*E.*) in children was recently estimated at 16% (10). Gut colonisation is a known risk factor for BSIs (11) and two-thirds of Tanzanian infants were faecal carriers (12). International guidelines advise to adapt recommended regimens for Gram-negative neonatal sepsis based on local pathogen susceptibility data (13). However, limited infrastructure and expertise in bacterial identification and antimicrobial susceptibility testing (AST) in Lower-Middle-Income-Countries like Tanzania can lead clinicians to administer non-optimal antibiotics (14), or lead patients to procure treatments from unlicensed vendors (15), further driving resistance. In addition, a key antibiotic used for AMR infections, ceftriaxone, has contraindications for neonatal sepsis use due to risk of hyperbilirubinemia (16) and calcium precipitate formation (17), leading to increased reliance on cefotaxime when ampicillin-gentamicin treatments fail (18). Therefore, further understanding of ARG amplification and their effect on AMR can improve genomic surveillance and treatment regimens. Here we describe the 18-week *in vivo* evolution of a community-acquired MDR ESBL-*E*, which exhibited de-amplification of an IS*26*-associated tandem ARG array.

## Materials and Methods

### Participant recruitment and sample collection

The *E. coli*, 1532WA and 1532MA, were isolated from a healthy infant, at age 6-weeks and at 6-months respectively. The individual was recruited for the ProRIDE probiotic placebo-controlled randomised clinical trial (19) set in Haydom Lutheran Hospital, Tanzania. The trial compared giving probiotics, Labinic (Biofloratech, Surrey, UK), or placebo for four weeks from birth, and stool samples were obtained at 6-weeks and at 6-months after birth. The subject infant was randomly designated into the control group who received the placebo. The infant was born at home with no birth complications, and no antibiotic usage or hospitalisations reported throughout the observation period.

### Species identification and antimicrobial susceptibility testing

Stool samples were collected and transferred into eSwab tubes (Copan Diagnostics, CA, USA) and stored at -80 °C at Haydom Lutheran Hospital. Then they were shipped on dry ice to Stavanger University Hospital, Norway and screened for ESBL-*E*. Briefly, eSwab samples were cultured on ChromID ESBL chromogenic selective agar (BioMerieux, Marcy l’Etoile, France) and non-selective blood agar for growth control. Plates were incubated at 35 °C for 18 – 24 h. Suspected ESBL-*E* colonies were identified to the species-level using MALDI-TOF (Bruker Daltonik, Bremen, Germany). AST was performed using the EUCAST disc diffusion method. Phenotypic ESBL-production was confirmed with the double disk approximation test (Becton Dickinson).

The two strains,1532WA and 1532MA, included in this study were identified as ESBL-*E. coli* and stored in Microbank cryovial system (Pro-lab Diagnostics, TX, USA) at -80 °C until transfer. They were then shipped to the Liverpool School of Tropical Medicine, UK and cultured in Luria-Bertani (LB, Sigma-Aldritch, MO, USA) agar plates for 18 – 24 h upon receipt. Streaks were suspended in 10% glycerol LB broth and stored at -70 °C.

### DNA extraction and whole-genome sequencing

Both strains were whole-genome-sequenced (WGS) on Illumina NovaSeq 6000 (CA, USA) and ONT MinION (Oxford Nanopore Technologies, Oxford, UK) platforms. Prior to sequencing, they were first cultured on LB agar and then LB broth, both supplemented with 2 µg/mL ceftriaxone (Sigma-Aldritch, MO, USA), to maintain ESBL selection pressure. The 6-months strain, 1532MA, was processed by using half a colony per sequencing platform to minimise inter-colony variation. The 6-weeks strain, 1532WA, could not be processed this way as it was Illumina-sequenced months prior to receipt of the 6-month isolate.

Isolates were submitted to MicrobesNG (www.microbesng.com), according to their strain submission guidelines, for Illumina WGS, 2 x 250 bp paired-end reads.

DNA extraction for ONT sequencing was performed using the MasterPure Complete DNA and RNA purification kit (LGC Biosearch Technologies, Hoddesdon, UK). Extracted DNA was quantified with a Qubit 4 fluorometer, following the dsDNA Broad-Range kit (Invitrogen, MA, USA) protocol, to ensure high yield. Concentration, integrity and length were confirmed on the Agilent Tapestation (Agilent, CA, USA) using the Genomic DNA ScreenTape. Samples were stored at 4 °C in DNA LoBind tubes (Eppendorf, Hamburg, Germany) after extraction and between ONT sequencing phases thereafter to minimise freeze-thaw shearing.

Extracts were barcoded using the Native Barcoding Kit (SQK-NBD114.24). DNA repair, end-prep, ligation and clean-up were performed, using the NEBNext Companion Module (NEB, MA, USA), according to manufacturer’s instructions.

R.10.4.1 flow cells were primed, with Bovine Serum Albumin (Invitrogen, MA, USA) in the priming mix, and loaded with the samples. Sequencing was run by MinKNOW v24.02.8 with default settings and duplex base-calling for 72 hours to produce FAST5 outputs. Base-calling and de-multiplexing were done using Guppy’s Super Accurate Simplex Model v1.1 (20).

### Quality control

Raw Illumina reads were manually assessed in FastQC v.0.11.5 (21) and trimmed using Trimmomatic v.0.39 (22) employing a sliding window cutoff of Q20 and removing adapter sequences. ONT reads were screened during the hybrid assembly pipeline.

### Assembly and annotation

Illumina and ONT reads were processed with the Hybracter v0.8.0 pipeline (23) with default settings to generate quality-controlled hybrid assemblies. In both genomes, two contigs were assembled, a complete 133 kbp plasmid and a > 5.31 Mbp chromosomal contig. The chromosomal contig was complete in 1532MA but was not in 1532WA.

Assemblies were manually assessed for quality using Quast v5.3.0 (24), then annotated against the full Bakta v1.10.4 database (25).

### Gene and plasmid replicon prediction

ARGs and plasmid replicons were predicted using ARIBA v2.11.1 (26) by querying *de novo* assembled Illumina reads against the SRST2-ARGANNOT (27,28) and PlasmidFinder (29) databases, respectively. These findings were cross-checked with Abricate v1.0.1 (30), using Hybracter assemblies.

Coverage and identity thresholds were respectively set at 80% and 90% for ARGs, and 60% and 95% for plasmid replicons. Predicted features satisfying these conditions were considered present. There were no differences in reported ARGs nor plasmid replicons between ARIBA and Abricate outputs.

An IS*26* reference sequence and the single-copy gene *rpoB* were added to the queried SRST2-ARGANNOT database. Achtmann-scheme sequence types were similarly called against the PubMLST database (31).

### Gene copy number estimation

The Bakta-annotated hybrid assembly of 1532WA revealed the presence of a terminal, chromosomal, repeating MDR transposon, which existed as a single copy in 1532MA. ONT reads were mapped back to the assembly with samtools v1.9 (32) to quantify coverage across the repetitive region and the genome. Illumina read coverage and count across ARGs were also compared against those of *rpoB,* using ARIBA, to estimate gene copy number. An adjustment was also applied to the read count according to gene size. These were plotted using ggplot2 (33) in R v4.5.0 (34).

### Average nucleotide identity and variant calling

Average nucleotide identity was calculated by FastANI v1.33 (35). SNPs were called using Breseq v0.38.3 (36), with default parameters, by mapping 1532MA’s Illumina reads onto 1532WA’s hybrid assembly.

### Growth curve analysis

Strains were cultured for 18 – 24 h in LB agar supplemented with 2 μg/mL ceftriaxone. A single colony was used to inoculate 10 mL LB broth and incubated at 37°C, shaking at 200 rpm, for 18 – 24 hours.

1950 μL of fresh LB broth was inoculated with 50 μL of broth culture to approximately 0.1 OD_600nm_, then measured on a spectrophotometer (Jenway, IL, US). A 1 μL aliquot was further diluted in LB broth to approximately 0.00002 OD_600nm_. 100 μL of the final dilution was added to 100 μL LB in each microplate well. Microplates were incubated at 37 °C for 24 hours in a SPECTROstar Omega automated microplate reader (BMG LabTech, Ortenberg, Germany). This experiment was repeated in triplicate for each strain.

The areas under the growth curves (AUCs) were calculated using the pracma R package (37). Due to the low sample size (n = 3) of repeats, non-normality was assumed, so a Kruskal-Wallis test for significance was applied.

### Antimicrobial susceptibility testing through disk diffusion

Phenotypic antimicrobial susceptibility testing was conducted following the European Committee on Antimicrobial Susceptibility Testing (EUCAST) guidelines. Briefly, strains were incubated in LB agar plates for 18 – 24 h. Two to three morphologically similar colonies were picked with a loop, dipped into 3 mL PBS and vortexed. A 0.5 MacFarland standard was prepared; confirming 0.08 – 0.13 OD_625nm_. This preparation was spread on Muller-Hinton agar (Sigma-Aldritch, MO, US).

Strains were tested against disks (OXOID, Basingstoke, UK) of ciprofloxacin (5 μg), gentamicin (30 μg), meropenem (10 μg), piperacillin-tazobactam (TZP, 110 μg), tetracycline (30 μg) and chloramphenicol (30 μg) and incubated at 37 °C for 18 h.

### Data availability

Read and assembly quality metrics, and scripts are available on GitHub in the repository: https://github.com/ralfhpulmones/deamplification. Assemblies are also accessible under the project code PRJNA1335781. Additional figures and supporting information are available in the Supplementary material.

## Results

### 1532WA and 1532MA are in vivo clones

Initial assessment of a larger dataset identified *E. coli* 1532MA, from an infant’s 6-months sample, as closely related to *E. coli* 1532WA, from the 6-week sample of the same infant. Both had an identical sequence type, identical ARGs (presence or absence) and plasmid replicons (Figure 1). An average nucleotide identity of 99.9983%, above the proposed consensus cutoff of 99.9% for same-strain clonality (38), further supported this. Subsequent Breseq variant calling identified only seven chromosomal SNPs between the two (Table 1) leading us to consider the strains highly related.

**Figure 1.**
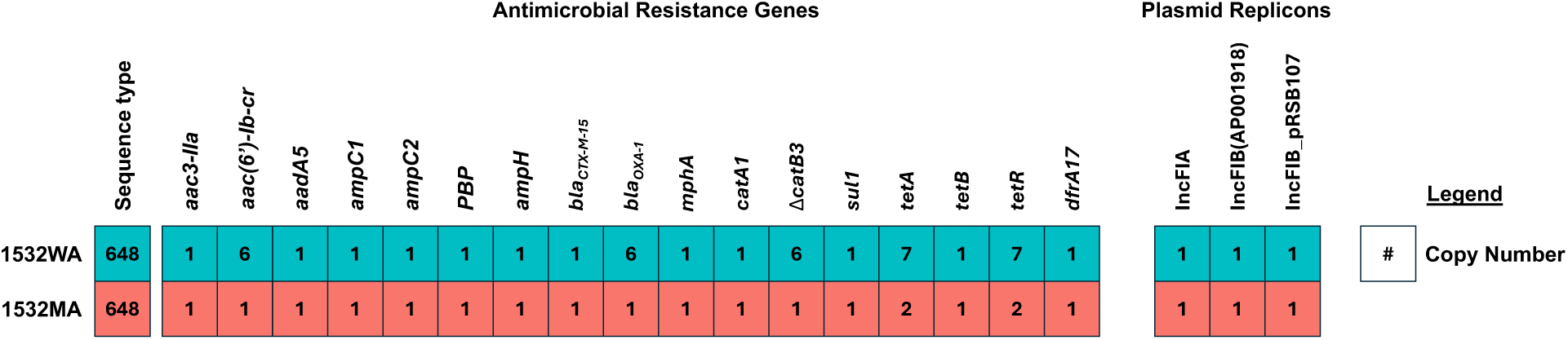
Predicted ARGs and plasmid replicons: ARGs and plasmid replicons predicted by ARIBA and cross-checked with Abricate. Copy numbers as annotated by Bakta and supported by Abricate.

**Table 1.**
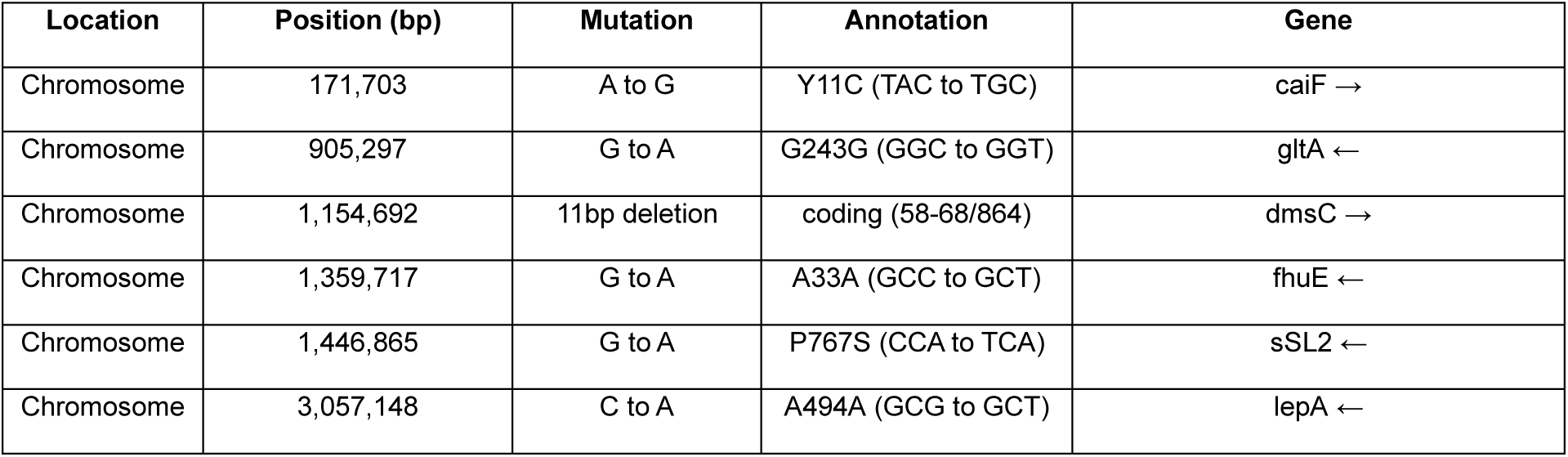
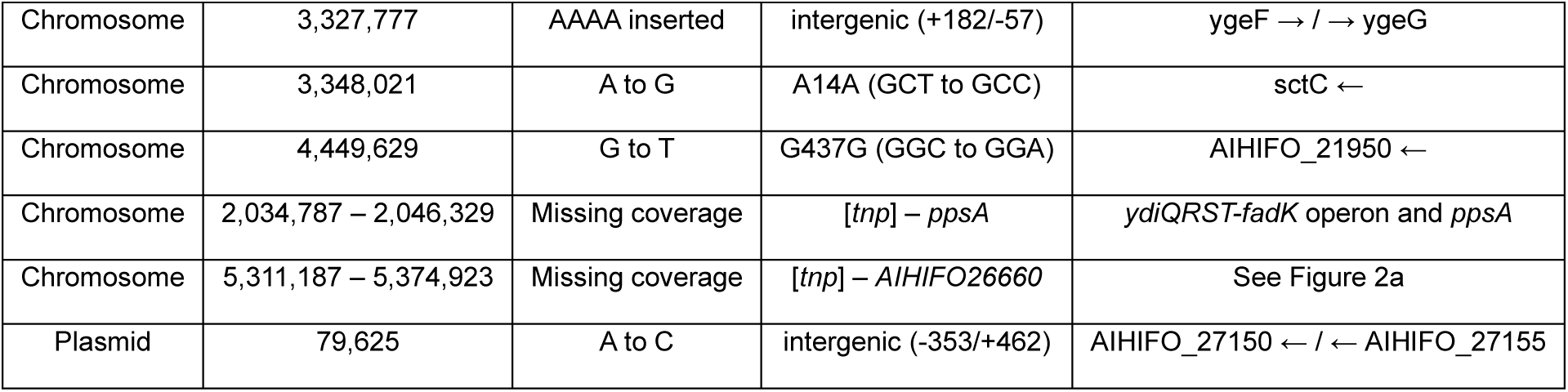
Breseq predicted mutations output: + indicates downstream gene, - indicates upstream. → is forward strand, ← is reverse strand. Includes approx. 63.5 kbp and 11.5 kbp regions in 1532WA which 1532MA’s reads did not cover, and were designated as “unassigned missing coverage evidence” separate from the predicted mutations.

| Location | Position (bp) | Mutation | Annotation | Gene |
| --- | --- | --- | --- | --- |
| Chromosome | 171,703 | A to G | Y11C (TAC to TGC) | caiF → |
| Chromosome | 905,297 | G to A | G243G (GGC to GGT) | gltA ← |
| Chromosome | 1,154,692 | 11bp deletion | coding (58-68/864) | dmsC → |
| Chromosome | 1,359,717 | G to A | A33A (GCC to GCT) | fhuE ← |
| Chromosome | 1,446,865 | G to A | P767S (CCA to TCA) | sSL2 ← |
| Chromosome | 3,057,148 | C to A | A494A (GCG to GCT) | lepA ← |
| Chromosome | 3,327,777 | AAAA inserted | intergenic (+182/-57) | ygeF → / → ygeG |
| Chromosome | 3,348,021 | A to G | A14A (GCT to GCC) | sctC ← |
| Chromosome | 4,449,629 | G to T | G437G (GGC to GGA) | AIHIFO_21950 ← |
| Chromosome | 2,034,787 – 2,046,329 | Missing coverage | [tnp] – ppsA | ydiQRST-fadK operon and ppsA |
| Chromosome | 5,311,187 – 5,374,923 | Missing coverage | [tnp] – AIHIFO26660 | See Figure 2a |
| Plasmid | 79,625 | A to C | intergenic (-353/+462) | AIHIFO_27150 ← / ← AIHIFO_27155 |

Mutations in *E. coli* 1532MA compared to *E. coli* 1532WA were distributed evenly across the chromosome with no hotspots, and none were linked to AMR but instead virulence or environmental adaptations, where known. Among the SNPs, only three were non-synonymous mutations, although none were associated with a known phenotype.

### Tandem array of multi-drug resistance transposon in ancestral week isolate

High quality assemblies of both strains were generated (N50 > 5,300,000 bp and L50 = 1), featuring two contigs each, one single-copy plasmid (132,802 bp) and their cognate chromosomes. 1532WA had an incomplete chromosomal contig which was 63,552 bp larger than the circularised chromosome of 1532MA. Although three plasmid replicons were predicted by ARIBA in both strains, all three mapped to the singular completed plasmid. Investigation into 1532WA’s annotated assembly revealed six tandem amplifications of a 9750 bp TU containing multiple ARGs, each with a flanking copy of IS*26*. These are terminally located on the assembled chromosome and is delimited by *yjhF*.

The TU consists of Tn*3-like*(*tnpA)*-*tetR-tetA-yedA-*ΔTn*1721*(*tnpA*)*-*IS*26*-*aac(6’)-Ib-cr-bla_OXA-1_-*Δ*catB3*-IS*26*, where Δ*catB3* is truncated by IS*26*, and this TU is part of a PCT. This PCT differs only in an additional flanking IS*26*, creating the structure IS*26*-Tn*3-like*(*tnpA)*-*tetR-tetA-yedA-*Tn*1721*(*tnpA*)*-*IS*26*-*aac(6’)-Ib-cr-bla_OXA-1_-*Δ*catB3*-IS*26* (Figure 2a). The ARGs within are likely to confer resistance to tetracyclines (*tetA*), aminoglycosides (*aac(6’)-Ib-cr*), fluoroquinolones (*aac(6’)-Ib-*cr) and penicillins (*bla_OXA-1_*) thus it can be classed as MDR conferring element. Strains with only this PCT should be susceptible to chloramphenicol due to the truncation of *catB3* (39). *yedA* encodes a putative transmembrane drug/metabolite transporter with unknown interactions with antimicrobials (40).

**Figure 2.**
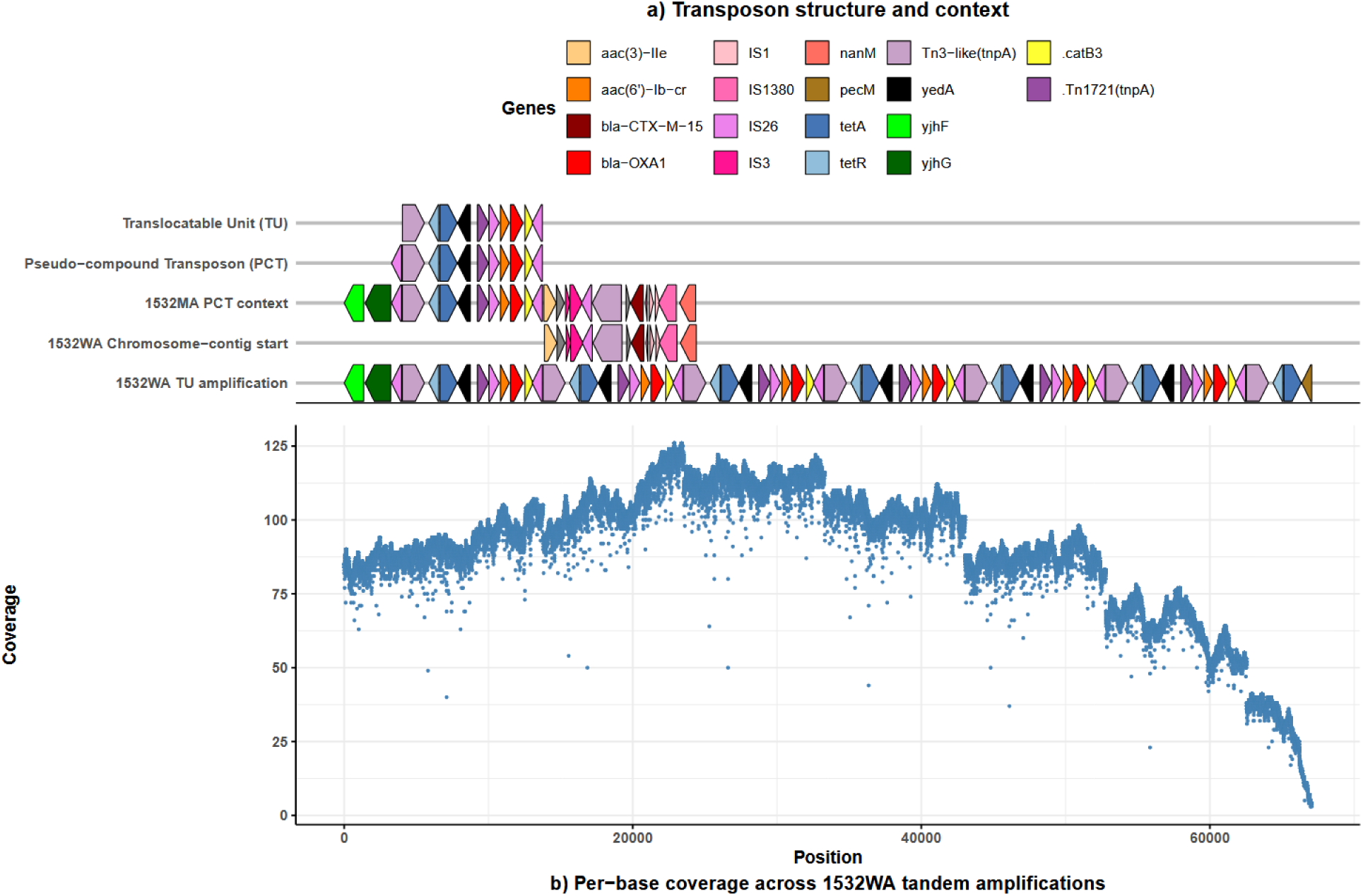
a Pseudo-Compound Transposon structure and context: Comparison of genomic structures around the MDR PCT in both strains. Orientation adjusted accordingly. **b Per-base coverage across 1532WA TU amplifications:** Coverage of ONT reads across the 1532WA TU amplifications. Aligned to the 1532WA TU amplification schematic above.

Only one PCT is present in 1532MA, therefore the remaining five TU repeats in 1532WA accounts for 48,750 bp of the 63,552 bp discrepancy. Meanwhile, a 4,664 bp incomplete TU, consisting of Tn*3*-like(*tnpA*)-*tetR*-*tetA*-*pecM*, capped the amplified region. The remaining 10 kbp is due to the absence of the approx. 11.5 kbp *ydiQRST-fadK* operon, alongside uncertainties in the boundaries of missing coverage regions.

The per-base ONT read coverage of the incomplete TU is considerably lower than elsewhere in the genome, and is also reduced to a lesser extent in the preceding PCT (Figure 2b). From the upstream *yjhF* to the terminal bases of the incomplete chromosomal contig of 1532WA, the mean per-base ONT read coverage was 90.1478 reads compared to 91.0183 mean reads for the entire chromosomal contig. Five TU repeats maintained coverage between 80 – 120 reads, while the sixth repeat had a mean coverage of approx. 60 reads across while the terminal bases of the contig had single-digit coverage. There is a peculiar pattern of lower coverage at two loci on all TU repeats. The first is in the intergenic region between Tn*3*-like(*tnpA*) and *tetR*, and the second was within *tetA.* This could be evidence of intra-clonal heterogeneity, with only some cells exhibiting these alleles, although this hypothesis and potential functional consequences to tetracycline resistance remain undetermined.

ARG copies were also estimated using Illumina read-mapping with ARIBA. Read counts and coverage across ARGs were compared to the single-copy *rpoB*. In 1532WA, ARGs present in the TU, including the repressor *tetR*, had generally elevated read counts (1.00 – 2.33x) and coverage (2.29 – 4.47x) compared to *rpoB*, and more than ARGs elsewhere in the genome. When adjusted for gene size, assuming no sequencing bias, mapped reads for TU ARGs were 6.16 – 13.66x greater than *rpoB. tetR* has consistently lower estimations, though its small size compared to other ARGs may make it more prone to natural read count variation. In contrast, IS*26* in 1532WA consistently had the most predicted copies due to the amplification of two IS*26* per TU and those present elsewhere in the genome. Overall, these values are concordant with ONT coverage and annotations.

**Figure 3.**
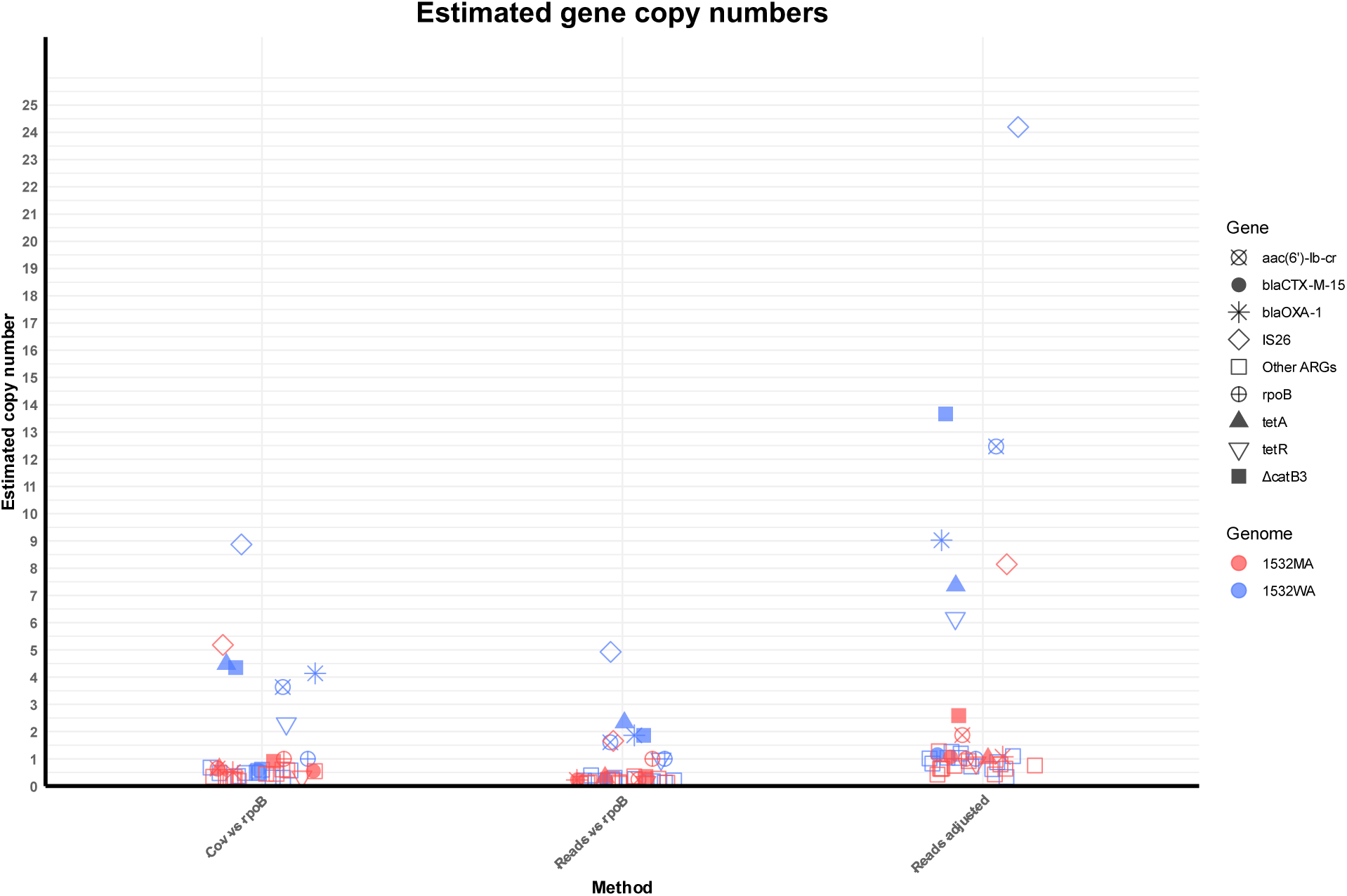
Estimated gene copy numbers: ARG and IS copy numbers estimated by Illumina read-mapping with ARIBA. ARGs inclusive to the TU apart from CTX-M which is included as a clinically relevant control outside of the TU. ARGs are highlighted as different shapes.

### Maintained fitness and resistance post de-amplification

Using AUC as a proxy for fitness, there was no fitness difference between 1532WA and 1532MA (p = 0.275). Therefore, TU de-amplification had no detectable effect on the fitness of 1532MA within the growth conditions of our experiments. Both strains exhibited phenotypic resistance to ciprofloxacin, chloramphenicol, tetracycline and gentamicin, and susceptibility to TZP and meropenem. Resistance to ciprofloxacin and tetracycline can be attributed to the PCT *aac(6’)-Ib-cr* and *tetA*, respectively. Immediately adjacent to the PCT in 1532MA is *aac*(*3*)*-IIa*, a resistance determinant to gentamicin, which is also present in 1532WA. Chloramphenicol resistance is unlikely to be from the Δ*catB3* (39) and instead the effect of a plasmid-borne *catA1*. No difference was observed in resistances to all tested agents except for gentamicin which 1532MA was more susceptible to than 1532WA.

**Figure 4.**
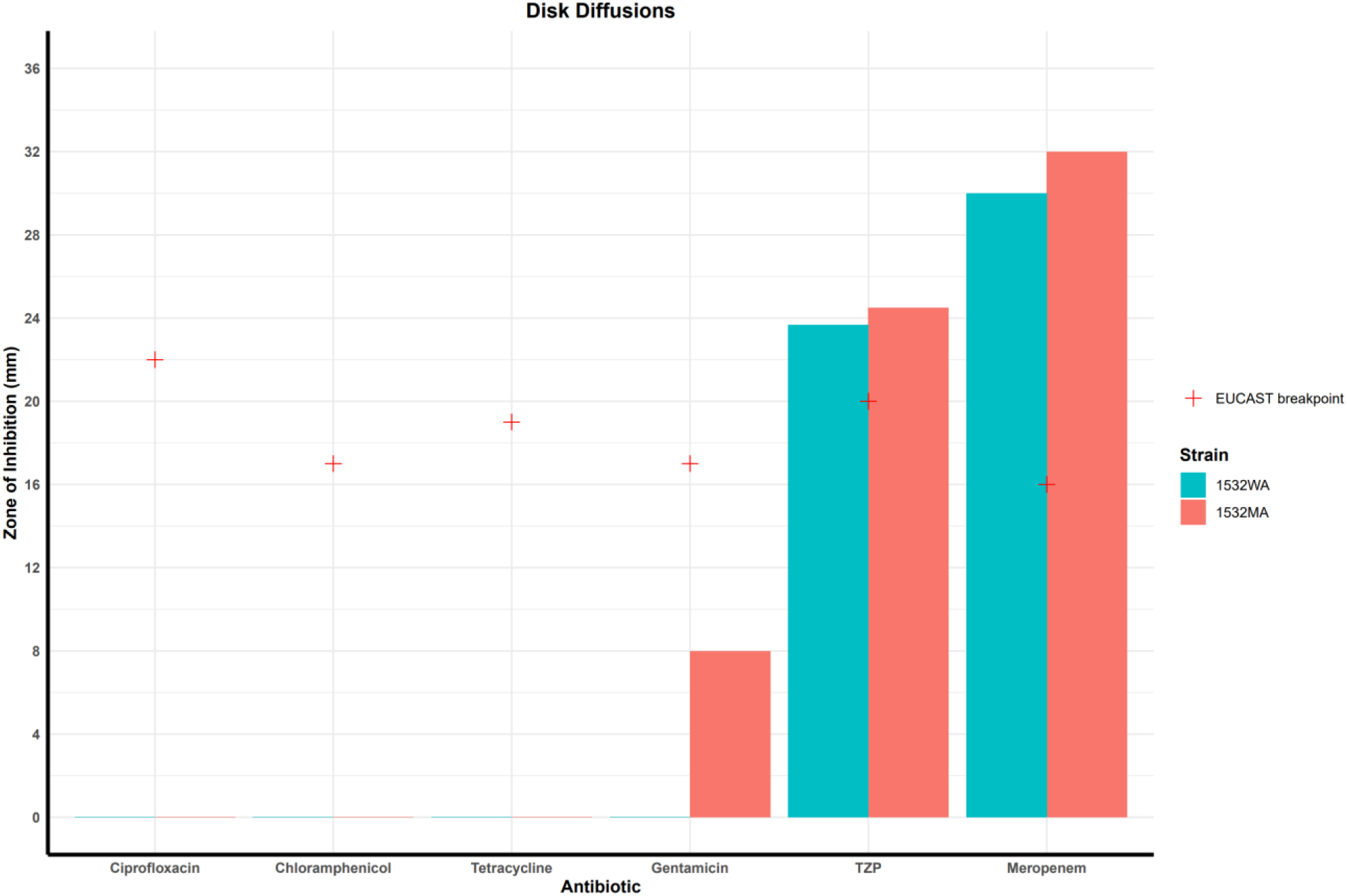
Disk diffusion: Conducted according to EUCAST guidelines and breakpoints.

*bla_OXA-1_* is a known contributor to modest TZP resistance (41), therefore it may be expected that more copies results in greater resistance, similar to *bla_TEM-1_* amplification (9,42,43). However, no significant increase in susceptibility to TZP was observed in 1532MA compared to 1532WA. Increased susceptibility to ciprofloxacin and tetracycline due to de-amplification could not be determined by disk diffusion as no zones of inhibition formed in either strain’s tests.

## Discussion

There is no consensus SNP threshold for clonality within human-colonising *E. coli*, but epidemiological studies often report evolution rates of 1.1 – 5.5 SNPs per genome per year (44,45) while long-term within-patient evolution studies (46,47) report 1 – 8 SNPs per genome per year. Meanwhile, a practical threshold of 5 – 17 SNPs is commonly used in outbreak analysis and real-time responses (48–50), with Public Health England initiating outbreak investigations if five isolates differing up to 5 SNPs are identified within 30 days (51). In contrast, a 100 SNPs threshold was proposed for One Health approaches to genomic surveillance (52).

These estimations were commonly derived from multi-centre studies across countries or used legacy technologies, comparing mainly adult populations. Evolution rates can also vary within even the same serotypes (53). Currently, there is no standardised methodology in sample source, collection, DNA extraction, sequencing, referencing, and variant calling (54–60) because of disparate research objectives and contexts.

Epidemiology utilises SNP analysis to, primarily, retrospectively track probable transmission pathways, and their thresholds are based on epidemiological linkage rather than laboratory data (48). Included in their estimates are SNPs generated in different hosts, and on intermediate surfaces and reservoirs, in contrast to within-patient studies where it is assumed that all SNPs are generated *in vivo*. However, it is currently impossible to track a cell and all its descendants outside of a sophisticated laboratory environment, thus there is an inherent uncertainty in the ancestry of any bacteria. Therefore, although a study may state clonality or ancestry between two strains, this is understood to mean that an extremely closely-related strain to one likely evolved to the other.

Our strains were isolated 18-weeks apart from the same infant, who was born at home with no record of hospitalisations or antibiotic use. Upon comparison they exhibited ten predicted mutations: two indels and eight SNPs, of which seven were chromosomal, alongside two large regions with missing coverage in 1532MA. This equates to 10.22 chromosomal SNPs per genome per year or 11.55 whole genome SNPs per genome per year. Therefore, using the previously mentioned criteria, our strains could be considered non-clonal; predicted to have accumulated approximately double the evolutionary distance expected within the 18 weeks between samplings.

However, our research contexts and those of the aforementioned studies are dissimilar, and we cannot assume that their SNPs estimates are generalisable to our strains. The infant microbiome is known to rapidly mature in the first months after birth. Initially, neonatal microbiomes are naïve upon vaginal birth and typically colonised by bacteria from their environment, their mother’s microbiome and breastmilk. As they transition to eating solid foods, typically around 5 months or earlier in some cultures (61), microbiome diversity generally increases (62). This increased diversity coincides with increased within-species genetic variation, and among *E. coli* this was estimated to be a 10-fold increase between birth and 5 - 6 months (63). It is probable that the introduction of novel microbial competitors, nutrition sources and the maturing immune system (64) promotes greater evolutionary pressure to adapt to new, more competitive niches in succession.

The brevity of the early-infancy life-stage is also incompatible with current research which typically covers multiple years, or decades, or evolution. Mutation accumulation is thought to approximate to a power-law model, in which bacteria introduced to a new environment relatively rapidly accumulate beneficial mutations but the rate decreases over time, approaching, but never reaching, zero (65). Strains under a multi-year observation period would, on average, exhibit a lower mutation rate than those observed more briefly. In a study of a recurrent urinary tract infection, 14 core genome SNPs were observed between *E. coli* isolated two years apart (3.5 SNPs per genome per year), while 16 SNPs differentiated the earlier strain versus another isolated a year later (8 SNPs per genome per year) (46). The power-law model alone could explain higher estimated mutation rates during short observation periods. Alongside the rapidly changing conditions in the infant microbiome, bacteria may be subjected to multiple rapid-adaptation periods during early infancy, resulting in higher SNP counts. Considering these factors and our data, there is sufficient evidence to assert that the community-acquired MDR strain 1532WA, or an extremely closely-related strain, likely persistently-colonised the infant and evolved into 1532MA.

An alternative hypothesis is that the infant was cleared of 1532WA naturally, and was colonised by 1532MA, which de-amplified the TU *ex vivo*, between 6-weeks and 6-months old. Indeed, in our parent study, most infants were no longer ESBL-*E* positive at 6-months old (19). This is a plausible scenario caused by limitations in tracking all similar longitudinal studies must accept.

It is also possible that both 1532WA and 1532MA were present at both timepoints but only one was isolated per timepoint due to our methodology. The strains may represent different lineages of a very recent common ancestor, which was not isolated, and in the 1532WA lineage the TU amplified whereas the 1532MA lineage retained the original structure, or vice versa. Indeed, a limitation of our study is that colony isolation was dependent on the uniqueness of the colony morphology rather than random isolation of multiple colonies per sample. Our method has a higher probability of isolating genetically distinct strains whereas the latter method is better for capturing the inter-colony variation within a sample (66), at the cost of greater redundancy. Due to the scale of the parent clinical trial and the cost of WGS, random isolation of multiple colonies was prohibitively expensive and much of the additional sequences would likely be from genetically identical colonies. We predict that intermediate strains with fewer SNP differences to either strain existed within our stool samples, and their mutations would show a generally cumulative pattern, however their recovery would be disproportionately expensive compared to their added resolution. Indeed, isolates are commonly referred to as clonal in contemporary literature without investigating inter-colony variation in the same sample (46).

1532WA and 1532MA differed the most in the presence of a tandem array of a TU in 1532WA which de-amplified into only the PCT in 1532MA. The TU structure (Tn*3-like*(*tnpA)*-*tetR-tetA-yedA-*ΔTn*1721*(*tnpA*)*-*IS*26*-*aac(6’)-Ib-cr-bla_OXA-1_-*Δ*catB3*-IS*26*) is similar to a Tn*1721*-like transposon reported in pEK499 (IS*26*-Δ*catB3*-*bla_OXA-1_*-*aacA4*-IS*26*-Tn*3*-like(*tnpA*)-*tetR*-*tetA*-*pecM*-ΔTn*1721*(*tnpA*)) (67), thus was also designated a Tn*1721*-like transposon. In our strains, downstream ΔTn*1721*(*tnpA*) is IS*26*-*aac(6’)-Ib-cr*-*bla_OXA-1_-*Δ*catB3*-IS*26*, with Δ*catB3*-IS*26* completing the PCT. In contrast, downstream pEK499’s ΔTn*1721*(*tnpA*) was Tn*501*(*tnpR*) and no nearby IS*26*. A PCT is formed in pEK499 but it excludes *tetR-tetA-pecM*, and instead includes *bla_CTX-M-15_*. ΔTn*1721*(*tnpA*) in pEK499 and our ΔTn*1721*(*tnpA*) were identical at all amino acids except in their flanks, likely due to their use of legacy sequencing technology. The transposons also differed in their carriage of *yedA* or *pecM*, however they are both uncharacterised members of the same drug/metabolite superfamily (68) and may be annotated as the other depending on coverage and the database used. Indeed, pEK499’s *pecM* and 1532WA’s TU *yedA* are identical while the terminal *pecM* in 1532WA shares 100% identity and 79% coverage with its *yedA*.

The amplified region spanned approximately 63 kbp and, unfortunately, there was insufficient ONT read coverage across the entire length. Therefore, the chromosome of 1532WA could not be resolved. However, independent analyses of the hybrid assembly, ONT reads and Illumina reads support the presence of at least five TUs, with a possible sixth. Furthermore, the incomplete TU is likely a mis-assembly of a few ONT reads. Its terminal gene was annotated as *pecM* by Bakta due to greater coverage when, in reality, it is a *yedA* which was truncated due to the mis-assembly. In all likelihood, a circularised chromosome assembly of 1532WA would show the amplifications adjacent to its chromosome-contig start, as seen in 1532MA (Figure 2a).

In hybrid assembly variant calling analysis, it is best practice to use the same colony for short-read and long-read WGS because it removes inter-colony variation, which can arise in pure isolate cultures because mutations can rapidly emerge. However, the practice remains uncommonly performed due to its practicality. This was done for 1532MA’s sequencing, but it was not possible for 1532WA because isolates were selected for additional ONT WGS based on whether the infant was ESBL-*E*. positive at both timepoints. Although this may be considered a limitation, ironically, it strengthens our findings. This is because the ONT and Illumina reads were from different colonies of the same isolate stock, and both support the presence of the TU amplifications independently of each other, and when hybrid assembled.

No discernible target-site duplications, deletions, truncations and inversions were observed, therefore amplification due to recombination is unlikely. Given the absence of these transposition-mechanism indicators and the PCT structure, IS*26*-mediated TCC is the most likely amplification mechanism (6). This mechanism requires two IS*26* to be directly-oriented, therefore it is unlikely that *bla_CTX-M-15_* can be amplified alongside the TU because there are no nearby IS*26* upstream *bla_CTX-M-15_* which satisfy this condition. However *aac*(*3*)*-IIa* could be amplified, alongside the defined TU or independently, because it has an upstream IS*26* with the same orientation as the TU IS*26*. Furthermore, the TU in 1532MA retains its potential for amplification as both directly-oriented IS*26* are intact.

Despite de-amplification, the strains could not be differentiated by phenotypic resistance to ciprofloxacin, chloramphenicol and tetracycline, as neither strain produced a zone of inhibition. Therefore, de-amplification did not increase susceptibility to an extent detectable by disk diffusion assay. Similarly, de-amplification of *bla_OXA-1_* did not result in a significant increase in susceptibility to TZP. Although gene copy number does not equate to increased expression, under selective pressure ARGs should be expressed. Therefore, it is unlikely that OXA-1 can hydrolyse piperacillin to clinical resistance, either because there were too few *bla_OXA-1_* amplifications to confer resistance, or OXA-1 affinity to piperacillin is too poor to be overcome by any increase in concentration.

De-amplification also coincided with greater susceptibility to gentamicin in 1532MA compared to 1532WA, and the cause is uncertain. The contribution of *yedA* to AMR is unknown and mutations identified by Breseq (Table 1) have no known associations to gentamicin susceptibility either. One copy of *aac(6’)-Ib-cr* is sufficient to confer resistance to aminoglycosides, through *N-*acetylation, except to gentamicin. However, amplification of another anti-aminoglycoside gene, *aphA1*, expanded resistance to tobramycin in *Acinetobacter baumannii* (8). Although *aphA1* acts through *O-*phosphorylation (69), both *aac(6’)-Ib-cr* and *aphA1* modify aminoglycosides at specific functional groups to disable binding to the 16s rRNA subunit A-site. Therefore, they may act similarly at high copies, e.g. enzyme saturation overcoming low affinity and efficiency. In *aphA1*, 65 amplifications extended resistance to tobramycin whereas *aac(6’)-Ib-cr* was amplified at least five times in 1532WA, and its de-amplification coincided with increased gentamicin susceptibility but not to clinical resistance. The potential for *aac(6’)-Ib-cr* amplifications to confer additional clinical resistances, at higher copies, is hitherto unexplored, and the dissemination of structures similar to our PCT could have future clinical consequences.

## Conclusion

This study identified a strain that featured tandem amplifications of a TU containing multiple ARGs, from an infant with no recorded hospitalisations or antibiotic usage. Its amplification was likely mediated by its encapsulating PCT, flanked by two directly-oriented copies of IS*26*, one of which truncated *catB3,* through TCC. Over 18-weeks *in vivo*, only the PCT remained despite no difference in fitness between the isolates. De-amplification of *bla_OXA-1_* led to no significant increase in susceptibility to TZP. In contrast, de-amplification coincided with increased susceptibility to gentamicin, possibly due to decreased copy number of *aac(6’)-Ib-cr*.

## References

1. WHO Bacterial Priority Pathogens List 2024: Bacterial Pathogens of Public Health Importance, to Guide Research, Development, and Strategies to Prevent and Control Antimicrobial Resistance. 1st ed. Geneva: World Health Organization; 2024. 1 p.

2. Tavakoli NP. Tipping the balance between replicative and simple transposition. The EMBO Journal. 2001 Jun 1;20(11):2923–30. doi:10.1093/emboj/20.11.2923

3. Munoz-Lopez M, Garcia-Perez J. DNA Transposons: Nature and Applications in Genomics. CG. 2010 Apr 1;11(2):115–28. doi:10.2174/138920210790886871

4. Nicolas E, Lambin M, Dandoy D, Galloy C, Nguyen N, Oger CA, et al. The Tn *3* -family of Replicative Transposons. Chandler M, Craig N, editors. Microbiol Spectr. 2015 Jul 2;3(4):3.4.14. doi:10.1128/microbiolspec.MDNA3-0060-2014

5. Harmer CJ, Hall RM. An analysis of the IS6/IS26 family of insertion sequences: is it a single family? Microbial Genomics. 2019 Sep 1;5(9). doi:10.1099/mgen.0.000291

6. Harmer CJ, Hall RM. IS26 and the IS26 family: versatile resistance gene movers and genome reorganizers. Microbiol Mol Biol Rev. 2024 Jun 27;88(2):e0011922. doi:10.1128/mmbr.00119-22 PubMed PMID: 38436262; PubMed Central PMCID: PMC11332343.

7. Harmer CJ, Moran RA, Hall RM. Movement of IS *26* -Associated Antibiotic Resistance Genes Occurs via a Translocatable Unit That Includes a Single IS *26* and Preferentially Inserts Adjacent to Another IS *26*. Bush K, editor. mBio. 2014 Oct 31;5(5):e01801–14. doi:10.1128/mBio.01801-14

8. Harmer CJ, Lebreton F, Stam J, McGann PT, Hall RM. Mechanisms of IS *26* -Mediated Amplification of the *aphA1* Gene Leading to Tobramycin Resistance in an Acinetobacter baumannii Isolate. Andam CP, editor. Microbiol Spectr. 2022 Oct 26;10(5):e02287–22. doi:10.1128/spectrum.02287-22

9. Hansen KH, Andreasen MR, Pedersen MS, Westh H, Jelsbak L, Schønning K. Resistance to piperacillin/tazobactam in Escherichia coli resulting from extensive IS26-associated gene amplification of blaTEM-1. Journal of Antimicrobial Chemotherapy. 2019 Nov 1;74(11):3179–83. doi:10.1093/jac/dkz349

10. Moyo SJ, Manyahi J, Aboud S, Mørch K, Roberts AP, Blomberg B, et al. Extended-spectrum-β-lactamase-producing Gram-negative bacteria are associated with high mortality in children with bloodstream infections in Dar es Salaam, Tanzania. BMC Infect Dis. 2025 Oct 27;25(1):1416. doi:10.1186/s12879-025-11629-4

11. Denis B, Lafaurie M, Donay JL, Fontaine JP, Oksenhendler E, Raffoux E, et al. Prevalence, risk factors, and impact on clinical outcome of extended-spectrum beta-lactamase-producing Escherichia coli bacteraemia: a five-year study. International Journal of Infectious Diseases. 2015 Oct;39:1–6. doi:10.1016/j.ijid.2015.07.010

12. Tellevik MG, Blomberg B, Kommedal Ø, Maselle SY, Langeland N, Moyo SJ. High Prevalence of Faecal Carriage of ESBL-Producing Enterobacteriaceae among Children in Dar es Salaam, Tanzania. Butaye P, editor. PLoS ONE. 2016 Dec 9;11(12):e0168024. doi:10.1371/journal.pone.0168024

13. Fuchs A, Bielicki J, Mathur S, Sharland M, Van Den Anker JN. Reviewing the WHO guidelines for antibiotic use for sepsis in neonates and children. Paediatrics and International Child Health. 2018 Dec 21;38(sup1):S3–15. doi:10.1080/20469047.2017.1408738

14. Styczynski A, Amin MB, Hoque KI, Parveen S, Md Pervez AF, Zeba D, et al. Perinatal colonization with extended-spectrum beta-lactamase-producing and carbapenem-resistant Gram-negative bacteria: a hospital-based cohort study. Antimicrob Resist Infect Control. 2024 Jan 29;13(1):13. doi:10.1186/s13756-024-01366-9

15. Belachew SA, Hall L, Selvey LA. Non-prescription dispensing of antibiotic agents among community drug retail outlets in Sub-Saharan African countries: a systematic review and meta-analysis. Antimicrob Resist Infect Control. 2021 Dec;10(1):13. doi:10.1186/s13756-020-00880-w

16. Martin E, Fanconi A, Kälin P, Zwingelstein C, Crevoisier C, Ruch W, et al. Ceftriaxone-bilirubin-albumin interactions in the neonate: An in vivo study. Eur J Pediatr. 1993 Jun;152(6):530–4. doi:10.1007/BF01955067

17. Steadman E, Raisch DW, Bennett CL, Esterly JS, Becker T, Postelnick M, et al. Evaluation of a Potential Clinical Interaction between Ceftriaxone and Calcium. Antimicrob Agents Chemother. 2010 Apr;54(4):1534–40. doi:10.1128/AAC.01111-09

18. Ministry of Health, Community Development, Gender, Eldery and Children. Standard treatment guidelines and national essential medicines list for Tanzania mainland [Internet]. The United Republic of Tanzania; 2021 [cited 2025 Oct 20]. Available from: https://www.moh.go.tz/storage/app/uploads/public/663/c8f/ceb/663c8fceb418d132695047.pdf

19. Klingenberg C, Justine M, Moyo SJ, Löhr IH, Gideon J, Mdoe P, et al. Home administration of a multistrain probiotic once per day for 4 weeks to newborn infants in Tanzania (ProRIDE): a double-blind, placebo-controlled randomised trial. The Lancet Global Health. 2025 Jun;13(6):e1082–90. doi:10.1016/S2214-109X(25)00064-6

20. Wick RR, Judd LM, Holt KE. Performance of neural network basecalling tools for Oxford Nanopore sequencing. Genome Biol. 2019 Dec;20(1):129. doi:10.1186/s13059-019-1727-y

21. Andrews S. FastQC [Internet]. 2010. (FastQC: A Quality Control Tool for High Throughput Sequence Data). Available from: https://www.bioinformatics.babraham.ac.uk/projects/fastqc/

22. Bolger AM, Lohse M, Usadel B. Trimmomatic: a flexible trimmer for Illumina sequence data. Bioinformatics. 2014 Aug 1;30(15):2114–20. doi:10.1093/bioinformatics/btu170

23. Bouras G, Houtak G, Wick RR, Mallawaarachchi V, Roach MJ, Papudeshi B, et al. Hybracter: enabling scalable, automated, complete and accurate bacterial genome assemblies. Microbial Genomics. 2024 May 8;10(5). doi:10.1099/mgen.0.001244

24. Gurevich A, Saveliev V, Vyahhi N, Tesler G. QUAST: quality assessment tool for genome assemblies. Bioinformatics. 2013 Apr 15;29(8):1072–5. doi:10.1093/bioinformatics/btt086

25. Schwengers O, Jelonek L, Dieckmann MA, Beyvers S, Blom J, Goesmann A. Bakta: rapid and standardized annotation of bacterial genomes via alignment-free sequence identification: Find out more about Bakta, the motivation, challenges and applications, here. Microbial Genomics. 2021 Nov 30;7(11). doi:10.1099/mgen.0.000685

26. Hunt M, Mather AE, Sánchez-Busó L, Page AJ, Parkhill J, Keane JA, et al. ARIBA: rapid antimicrobial resistance genotyping directly from sequencing reads. Microbial Genomics. 2017 Oct 1;3(10). doi:10.1099/mgen.0.000131

27. Inouye M, Dashnow H, Raven LA, Schultz MB, Pope BJ, Tomita T, et al. SRST2: Rapid genomic surveillance for public health and hospital microbiology labs. Genome Med. 2014 Nov 20;6(11):90. doi:10.1186/s13073-014-0090-6

28. Gupta SK, Padmanabhan BR, Diene SM, Lopez-Rojas R, Kempf M, Landraud L, et al. ARG-ANNOT, a New Bioinformatic Tool To Discover Antibiotic Resistance Genes in Bacterial Genomes. Antimicrob Agents Chemother. 2014 Jan;58(1):212–20. doi:10.1128/AAC.01310-13

29. Carattoli A, Zankari E, García-Fernández A, Voldby Larsen M, Lund O, Villa L, et al. *In Silico* Detection and Typing of Plasmids using PlasmidFinder and Plasmid Multilocus Sequence Typing. Antimicrob Agents Chemother. 2014 Jul;58(7):3895–903. doi:10.1128/AAC.02412-14

30. T. Seemann. ABRicate [Internet]. 2020. Available from: https://github.com/tseemann/abricate.

31. Jolley KA, Bray JE, Maiden MCJ. Open-access bacterial population genomics: BIGSdb software, the PubMLST.org website and their applications. Wellcome Open Res. 2018 Sep 24;3:124. doi:10.12688/wellcomeopenres.14826.1

32. Li H, Handsaker B, Wysoker A, Fennell T, Ruan J, Homer N, et al. The Sequence Alignment/Map format and SAMtools. Bioinformatics. 2009 Aug 15;25(16):2078–9. doi:10.1093/bioinformatics/btp352

33. Wickham H, Averick M, Bryan J, Chang W, McGowan L, François R, et al. Welcome to the Tidyverse. JOSS. 2019 Nov 21;4(43):1686. doi:10.21105/joss.01686

34. R Core Team [Internet]. R Foundation for Statistical Computing, Vienna, Austria.; 2025. (R: A Language and Environment for Statistical Computing). Available from: https://www.R-project.org

35. Jain C, Rodriguez-R LM, Phillippy AM, Konstantinidis KT, Aluru S. High throughput ANI analysis of 90K prokaryotic genomes reveals clear species boundaries. Nat Commun. 2018 Nov 30;9(1):5114. doi:10.1038/s41467-018-07641-9

36. Deatherage DE, Barrick JE. Identification of Mutations in Laboratory-Evolved Microbes from Next-Generation Sequencing Data Using breseq. In: Sun L, Shou W, editors. Engineering and Analyzing Multicellular Systems [Internet]. New York, NY: Springer New York; 2014 [cited 2025 Jun 10]. p. 165–88. (Methods in Molecular Biology). Available from: https://link.springer.com/10.1007/978-1-4939-0554-6_12 doi:10.1007/978-1-4939-0554-6_12

37. pracma [Internet]. 2023. (pracma: Practical Numerical Math Functions). Available from: https://CRAN.R-project.org/package=pracma

38. Rodriguez-R LM, Conrad RE, Viver T, Feistel DJ, Lindner BG, Venter SN, et al. An ANI gap within bacterial species that advances the definitions of intra-species units. Jouline IB, editor. mBio. 2024 Jan 16;15(1):e02696–23. doi:10.1128/mbio.02696-23

39. Graf FE, Goodman RN, Gallichan S, Forrest S, Picton-Barlow E, Fraser AJ, et al. Molecular mechanisms of re-emerging chloramphenicol susceptibility in extended-spectrum beta-lactamase-producing Enterobacterales. Nat Commun. 2024 Oct 18;15(1):9019. doi:10.1038/s41467-024-53391-2

40. Desai KK, Miller BG. Recruitment of genes and enzymes conferring resistance to the nonnatural toxin bromoacetate. Proc Natl Acad Sci USA. 2010 Oct 19;107(42):17968–73. doi:10.1073/pnas.1007559107

41. Livermore DM, Day M, Cleary P, Hopkins KL, Toleman MA, Wareham DW, et al. OXA-1 β-lactamase and non-susceptibility to penicillin/β-lactamase inhibitor combinations among ESBL-producing *Escherichia coli*. Journal of Antimicrobial Chemotherapy. 2019 Feb 1;74(2):326–33. doi:10.1093/jac/dky453

42. Schechter LM, Creely DP, Garner CD, Shortridge D, Nguyen H, Chen L, et al. Extensive Gene Amplification as a Mechanism for Piperacillin-Tazobactam Resistance in Escherichia coli. mBio. 2018 Apr 24;9(2):e00583–18. doi:10.1128/mBio.00583-18 PubMed PMID: 29691340; PubMed Central PMCID: PMC5915731.

43. Hubbard ATM, Mason J, Roberts P, Parry CM, Corless C, van Aartsen J, et al. Piperacillin/tazobactam resistance in a clinical isolate of Escherichia coli due to IS26-mediated amplification of blaTEM-1B. Nat Commun. 2020 Oct 1;11(1):4915. doi:10.1038/s41467-020-18668-2 PubMed PMID: 33004811; PubMed Central PMCID: PMC7530762.

44. Von Mentzer A, Connor TR, Wieler LH, Semmler T, Iguchi A, Thomson NR, et al. Identification of enterotoxigenic Escherichia coli (ETEC) clades with long-term global distribution. Nat Genet. 2014 Dec;46(12):1321–6. doi:10.1038/ng.3145

45. White RT, Bull MJ, Barker CR, Arnott JM, Wootton M, Jones LS, et al. Genomic epidemiology reveals geographical clustering of multidrug-resistant Escherichia coli ST131 associated with bacteraemia in Wales. Nat Commun. 2024 Feb 14;15(1):1371. doi:10.1038/s41467-024-45608-1

46. Forde BM, Roberts LW, Phan MD, Peters KM, Fleming BA, Russell CW, et al. Population dynamics of an Escherichia coli ST131 lineage during recurrent urinary tract infection. Nat Commun. 2019 Aug 13;10(1):3643. doi:10.1038/s41467-019-11571-5

47. Parsons JB, Sidders AE, Velez AZ, Hanson BM, Angeles-Solano M, Ruffin F, et al. In-patient evolution of a high-persister *Escherichia coli* strain with reduced in vivo antibiotic susceptibility. Proc Natl Acad Sci USA. 2024 Jan 16;121(3):e2314514121. doi:10.1073/pnas.2314514121

48. Ludden C, Coll F, Gouliouris T, Restif O, Blane B, Blackwell GA, et al. Defining nosocomial transmission of Escherichia coli and antimicrobial resistance genes: a genomic surveillance study. The Lancet Microbe. 2021 Sep;2(9):e472–80. doi:10.1016/S2666-5247(21)00117-8

49. Mills EG, Martin MJ, Luo TL, Ong AC, Maybank R, Corey BW, et al. A one-year genomic investigation of Escherichia coli epidemiology and nosocomial spread at a large US healthcare network. Genome Med. 2022 Dec 30;14(1):147. doi:10.1186/s13073-022-01150-7

50. Dallman TJ, Greig DR, Gharbia SE, Jenkins C. Phylogenetic structure of Shiga toxin-producing Escherichia coli O157:H7 from sub-lineage to SNPs. Microbial Genomics. 2021 Mar 1;7(3). doi:10.1099/mgen.0.000544

51. Dallman TJ, Byrne L, Ashton PM, Cowley LA, Perry NT, Adak G, et al. Whole-Genome Sequencing for National Surveillance of Shiga Toxin–Producing *Escherichia coli* O157. Clin Infect Dis. 2015 Aug 1;61(3):305–12. doi:10.1093/cid/civ318

52. Watt AE, Cummins ML, Donato CM, Wirth W, Porter AF, Andersson P, et al. Parameters for one health genomic surveillance of Escherichia coli from Australia. Nat Commun. 2025 Jan 2;16(1):17. doi:10.1038/s41467-024-55103-2

53. Weinroth MD, Clawson ML, Arthur TM, Wells JE, Brichta-Harhay DM, Strachan N, et al. Rates of evolutionary change of resident Escherichia coli O157:H7 differ within the same ecological niche. BMC Genomics. 2022 Apr 7;23(1):275. doi:10.1186/s12864-022-08497-6

54. Holmes A, Allison L, Ward M, Dallman TJ, Clark R, Fawkes A, et al. Utility of Whole-Genome Sequencing of Escherichia coli O157 for Outbreak Detection and Epidemiological Surveillance. Diekema DJ, editor. J Clin Microbiol. 2015 Nov;53(11):3565–73. doi:10.1128/JCM.01066-15

55. Joensen KG, Scheutz F, Lund O, Hasman H, Kaas RS, Nielsen EM, et al. Real-Time Whole-Genome Sequencing for Routine Typing, Surveillance, and Outbreak Detection of Verotoxigenic Escherichia coli. Carroll KC, editor. J Clin Microbiol. 2014 May;52(5):1501–10. doi:10.1128/JCM.03617-13

56. Dallman TJ, Ashton PM, Byrne L, Perry NT, Petrovska L, Ellis R, et al. Applying phylogenomics to understand the emergence of Shiga-toxin-producing Escherichia coli O157:H7 strains causing severe human disease in the UK. Microbial Genomics. 2015 Sep 14;1(3). doi:10.1099/mgen.0.000029

57. Roer L, Hansen F, Thomsen MCF, Knudsen JD, Hansen DS, Wang M, et al. WGS-based surveillance of third-generation cephalosporin-resistant Escherichia coli from bloodstream infections in Denmark. Journal of Antimicrobial Chemotherapy. 2017 Jul;72(7):1922–9. doi:10.1093/jac/dkx092

58. Hammerum AM, Hansen F, Nielsen HL, Jakobsen L, Stegger M, Andersen PS, et al. Use of WGS data for investigation of a long-term NDM-1-producing *Citrobacter freundii* outbreak and secondary *in vivo* spread of *bla*_NDM-1_ to *Escherichia coli*, *Klebsiella pneumoniae* and *Klebsiella oxytoca*. J Antimicrob Chemother. 2016 Nov;71(11):3117–24. doi:10.1093/jac/dkw289

59. Foster-Nyarko E, Alikhan NF, Ikumapayi UN, Sarwar G, Okoi C, Tientcheu PEM, et al. Genomic diversity of *Escherichia coli* from healthy children in rural Gambia. PeerJ. 2021 Jan 6;9:e10572. doi:10.7717/peerj.10572

60. Reeves PR, Liu B, Zhou Z, Li D, Guo D, Ren Y, et al. Rates of Mutation and Host Transmission for an Escherichia coli Clone over 3 Years. Lin B, editor. PLoS ONE. 2011 Oct 27;6(10):e26907. doi:10.1371/journal.pone.0026907

61. Van Dijk M, Hunnius S, Van Geert P. The dynamics of feeding during the introduction to solid food. Infant Behavior and Development. 2012 Apr;35(2):226–39. doi:10.1016/j.infbeh.2012.01.001

62. Yang I, Corwin EJ, Brennan PA, Jordan S, Murphy JR, Dunlop A. The Infant Microbiome: Implications for Infant Health and Neurocognitive Development. Nursing Research. 2016 Jan;65(1):76–88. doi:10.1097/NNR.0000000000000133

63. Chen DW, Garud NR. Rapid evolution and strain turnover in the infant gut microbiome. Genome Res. 2022 Jun;32(6):1124–36. doi:10.1101/gr.276306.121

64. Sjögren YM, Tomicic S, Lundberg A, Böttcher MF, Björkstén B, Sverremark-Ekström E, et al. Influence of early gut microbiota on the maturation of childhood mucosal and systemic immune responses: Gut microbiota and immune responses. Clin Experimental Allergy. 2009 Dec;39(12):1842–51. doi:10.1111/j.1365-2222.2009.03326.x

65. Lenski RE. Experimental evolution and the dynamics of adaptation and genome evolution in microbial populations. The ISME Journal. 2017 Oct 1;11(10):2181–94. doi:10.1038/ismej.2017.69

66. Gallichan S, Forrest S, Picton-Barlow E, McKeown C, Moore M, Heinz E, et al. Optimized methods for the targeted surveillance of extended-spectrum beta-lactamase-producing Escherichia coli in human stool. Microbiol Spectr. 2025 Jan 7;13(1):e0105824. doi:10.1128/spectrum.01058-24 PubMed PMID: 39576089; PubMed Central PMCID: PMC11705872.

67. Woodford N, Carattoli A, Karisik E, Underwood A, Ellington MJ, Livermore DM. Complete nucleotide sequences of plasmids pEK204, pEK499, and pEK516, encoding CTX-M enzymes in three major Escherichia coli lineages from the United Kingdom, all belonging to the international O25:H4-ST131 clone. Antimicrob Agents Chemother. 2009 Oct;53(10):4472–82. doi:10.1128/AAC.00688-09 PubMed PMID: 19687243; PubMed Central PMCID: PMC2764225.

68. Jack DL, Yang NM, H. Saier M. The drug/metabolite transporter superfamily. European Journal of Biochemistry. 2001 Jul;268(13):3620–39. doi:10.1046/j.1432-1327.2001.02265.x

69. Vetting MW, Park CH, Hegde SS, Jacoby GA, Hooper DC, Blanchard JS. Mechanistic and Structural Analysis of Aminoglycoside *N* -Acetyltransferase AAC(6′)-Ib and Its Bifunctional, Fluoroquinolone-Active AAC(6′)-Ib-cr Variant. Biochemistry. 2008 Sep 16;47(37):9825–35. doi:10.1021/bi800664x

