## Supplementary materials for "*In vivo* de-amplification of a multi-resistance pseudo-compound transposon in Escherichia coli"

**Supplementary material**

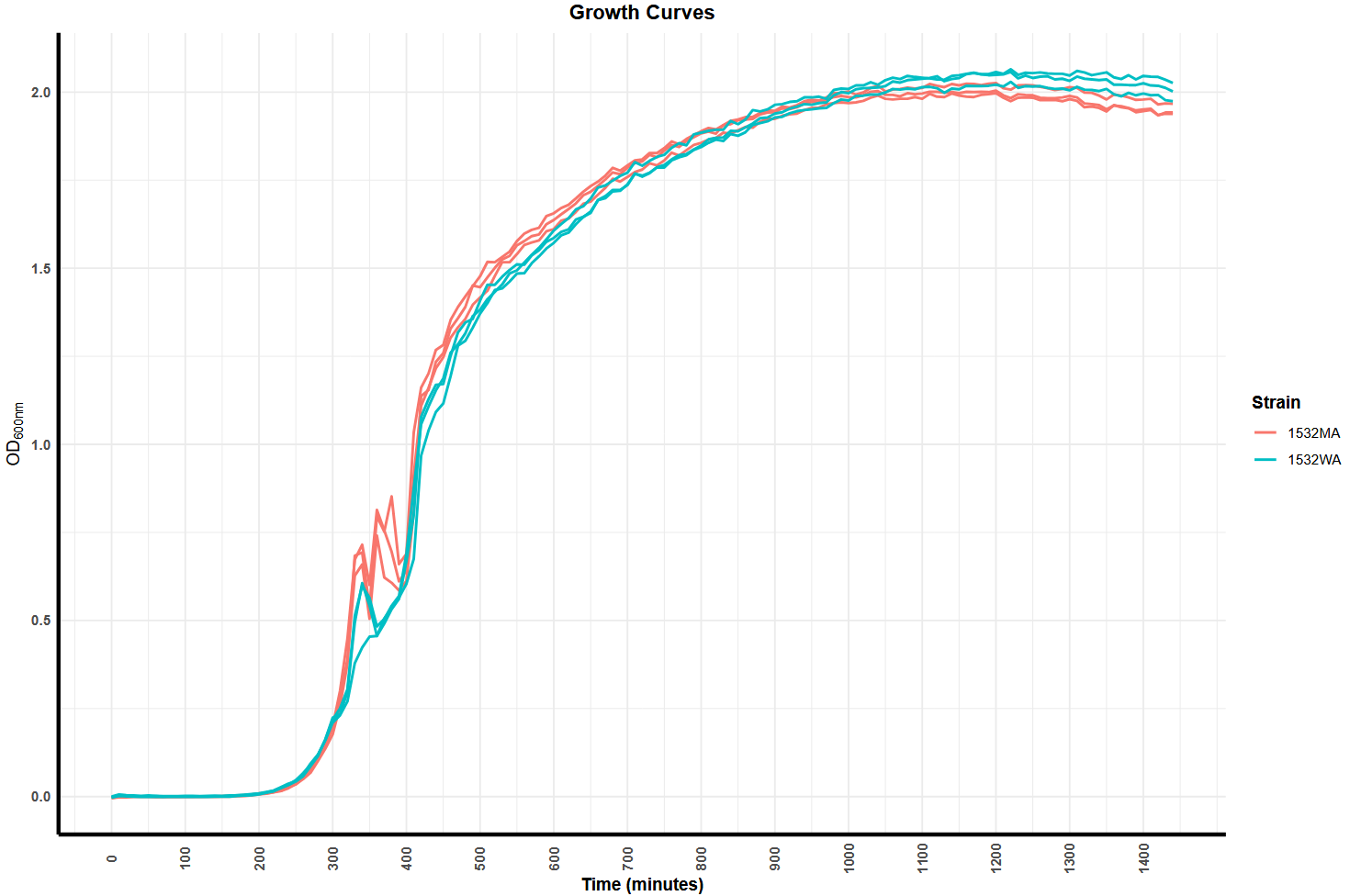

**Supplement 1: Strain growth curves:** OD_600nm_ from an inoculum of 0.00001 OD_600nm_ in 200 μL LB broth over 24 h

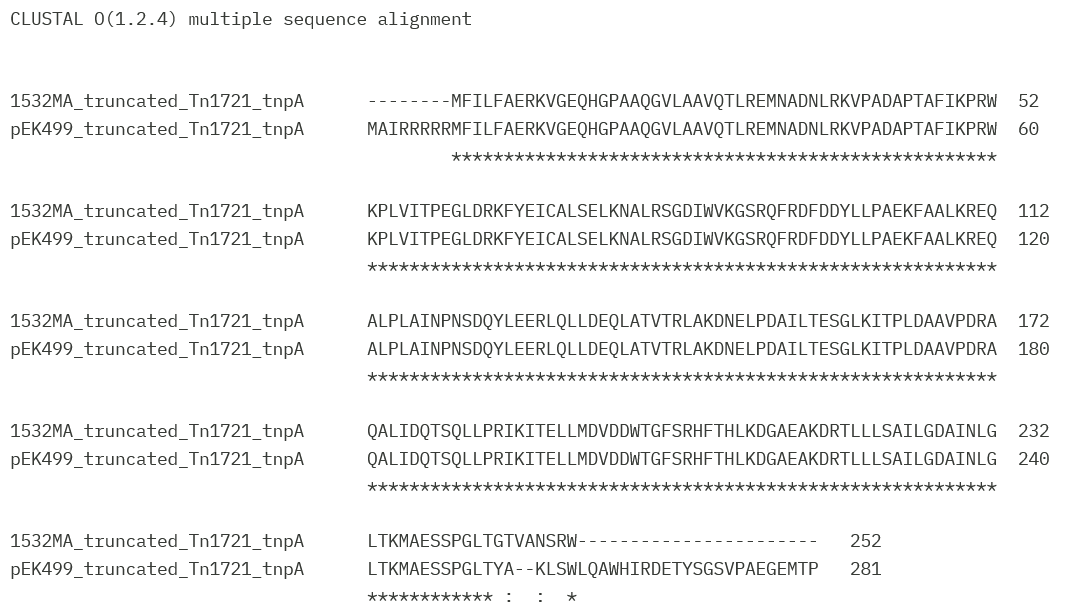

**Supplement 2: Clustal Omega alignment of pEK499 ΔTn1721 and 1532WA’s TU Tn3-like tnpA**

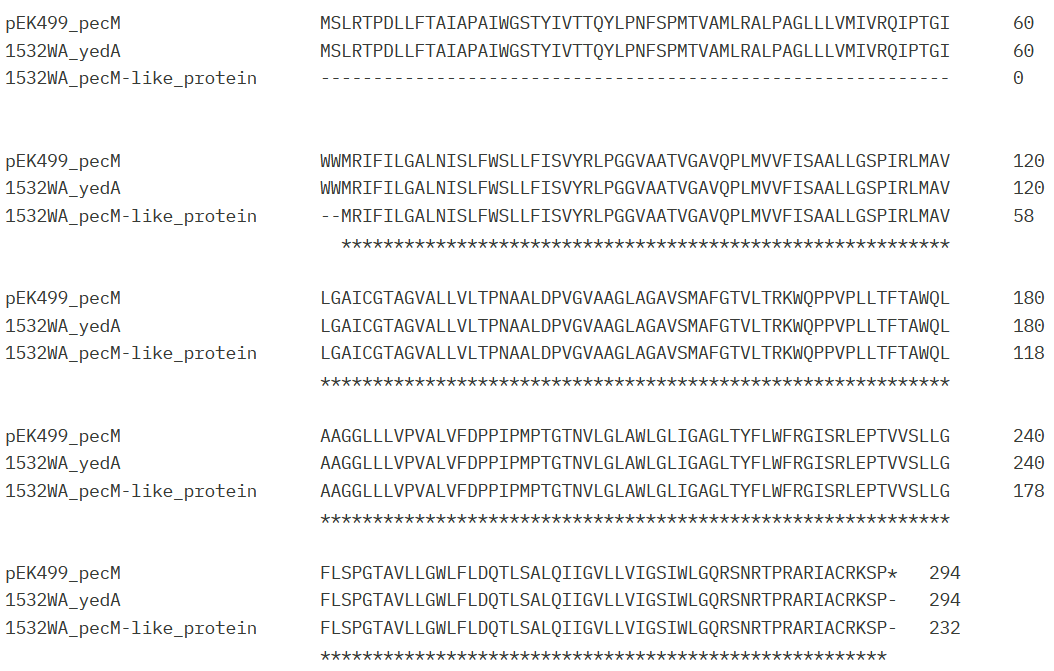

**Supplement 3: Clustal Omega alignment of pEK499 pecM and 1532WA’s TU yedA and pecM-like protein**

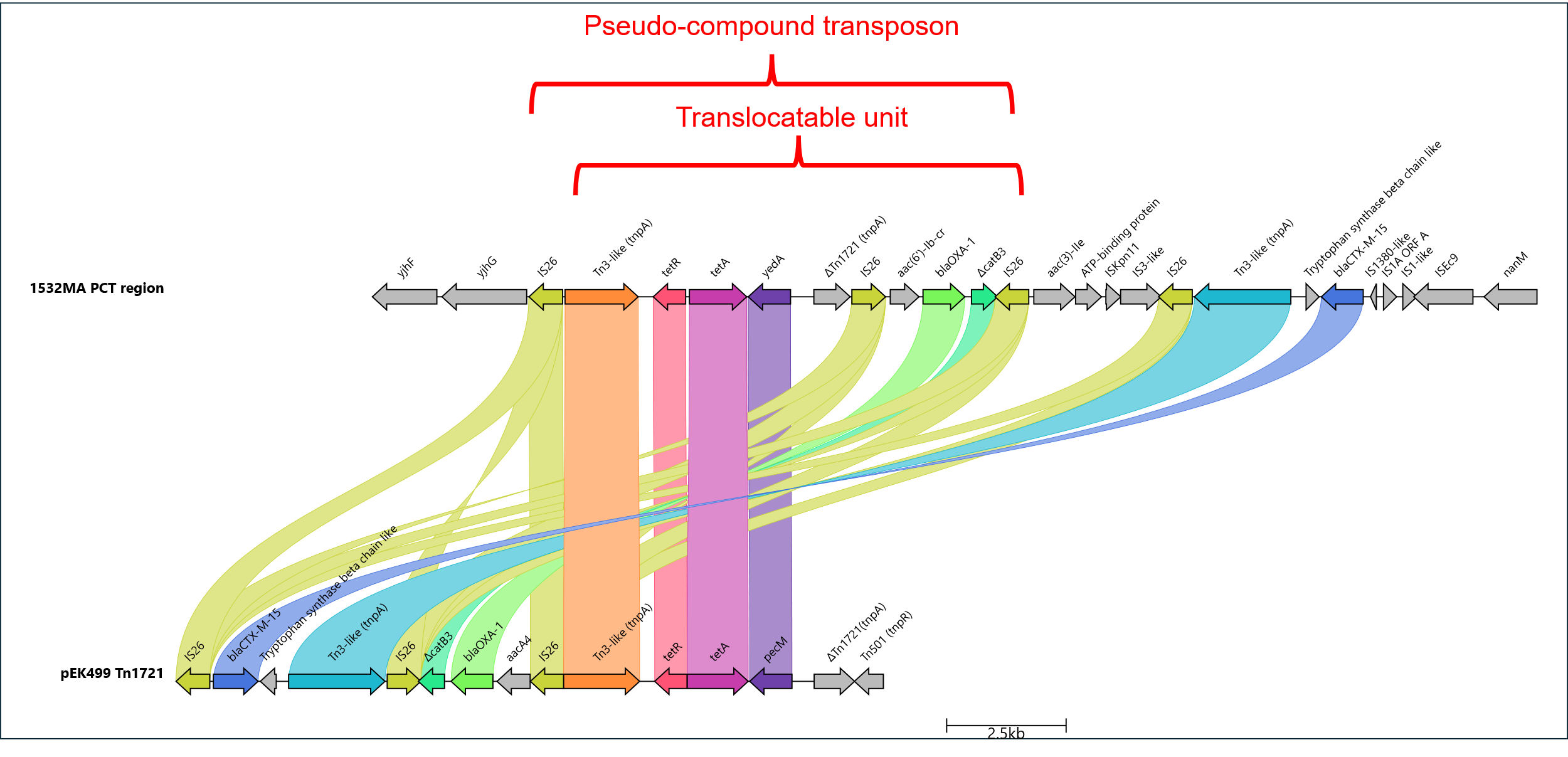

**Supplement 4: Clinker alignment of 1532MA PCT region and pEK499 Tn1721-like:** Manually re-named genes for consistency
